# Foxc1 establishes enhancer accessibility for craniofacial cartilage differentiation

**DOI:** 10.1101/2020.10.15.340703

**Authors:** Pengfei Xu, Haoze Vincent Yu, Kuo-Chang Tseng, Mackenzie Flath, Peter Fabian, Neil Segil, J. Gage Crump

## Abstract

The specification of cartilage requires Sox9, a transcription factor with broad roles for organogenesis outside the skeletal system. How Sox9 gains selective access to cartilage-specific cis-regulatory regions during skeletal development had remained unclear. By analyzing chromatin accessibility during the differentiation of neural crest cells into chondrocytes of the zebrafish head, we find that cartilage-associated chromatin accessibility is dynamically established. Cartilage-associated regions that become accessible after neural crest migration are co-enriched for Sox9 and Fox transcription factor binding motifs. In zebrafish lacking Foxc1 paralogs, we find a global decrease in chromatin accessibility in chondrocytes, consistent with a later loss of dorsal facial cartilages. Zebrafish transgenesis assays confirm that many of these Foxc1-dependent elements function as enhancers with region- and stage-specific activity in facial cartilages. We propose that Foxc1-dependent chromatin accessibility helps directs the versatile Sox9 protein to a chondrogenic program in the face.

**Highlights:**

- Dynamic chromatin accessibility across facial cartilage development
- Co-enrichment of Fox- and Sox-binding motifs in accessible regions
- Foxc1 establishes accessibility in a subset of facial cartilage enhancers
- Modular activity of Foxc1-dependent cartilage enhancers in zebrafish

## Introduction

Cartilage is the first skeletal type to be specified in the vertebrate body, providing important templates for later bone development and providing flexibility at joint surfaces and within the nose, ear, ribs, and larynx. The transcription factor Sox9 is essential for chondrogenic differentiation in all vertebrates examined (Lefebvre et al., 1997; Bi et al., 1999; Mori-Akiyama et al., 2003; Yan et al., 2005), yet it also has widespread roles outside the skeletal system, including in the reproductive system, kidney, liver, and skin (Jo et al., 2014). How Sox9 is directed to a chondrogenic program in trunk mesoderm and cranial neural crest-derived cells (CNCCs) has remained unclear. Sox9 is known to directly bind to a number of cis-regulatory elements adjacent to chondrogenic genes, including *Col2a1, Col10a1*, and *Acan* (Lefebvre et al., 1997; Dy et al., 2012; Askary et al., 2015; Ohba et al., 2015). However, it is not required for chromatin accessibility at these same elements (Liu et al., 2018), suggesting other unknown factors may first open chromatin at chondrogenic enhancers for later activation by Sox9.

Forkhead-domain (Fox) family transcription factors are excellent candidates for making chondrogenic enhancers accessible for later Sox9 activation. In the endoderm lineage, HNF3/FoxA binds closed chromatin at enhancers and makes these more accessible (Cirillo et al., 2002). A similar chromatin accessibility role for Foxd3 in the early neural crest lineage has also been proposed (Lukoseviciute et al., 2018). In mice, loss of *Foxc1* results in widespread cartilage and bone defects (Kume et al., 1998), including impaired tracheal and rib cartilages (Hong et al., 1999), loss of calvarial bone due to premature ossification (Rice et al., 2003; Vivatbutsiri et al., 2008; Sun et al., 2013), syngnathia (Inman et al., 2013), and disruption of endochondral bone maturation (Yoshida et al., 2015). In zebrafish, loss of both Foxc1 paralogs (*foxc1a* and *foxc1b*) results in severe reductions of dorsal cartilages of the upper face, which are preceded by reduced expression of several Sox9 targets, including *col2a1a, acana, matn1*, and *matn4* (Xu et al., 2018). Chromatin immunoprecipitation followed by deep sequencing (ChIP-Seq) using a Sox9 antibody in dissected mouse rib and nose cartilage revealed enrichment of Fox binding motifs within Sox9-bound cis-regulatory sequences near chondrogenic genes (Ohba et al., 2015). Here, we use profiling of chromatin accessibility in wild-type and mutant zebrafish facial cartilages and find that Foxc1 paralogs are required for accessibility and activity of a number of cartilage enhancers. These findings support a model by which Foxc1 restricts Sox9 activity to a chondrogenic program by establishing selective accessibility of cartilage enhancers in CNCC-derived mesenchyme.

## Results and Discussion

### Chromatin accessibility landscape in chondrocytes of the zebrafish face

In order to identify potential cis-regulatory elements important for facial cartilage development, we performed a genome-wide analysis of chromatin accessibility in chondrocytes from 72 hours post-fertilization (hpf) zebrafish. We labeled chondrocytes by co-expression of *sox10:Dsred* and *col2a1a:GFP* transgenes (Figure 1A) and isolated double-positive and control double-negative cells by fluorescence-activated cell sorting (FACS). We then subjected these cells to a modified “micro” version of the assay for transposase accessible chromatin followed by next-generation sequencing (μATACseq). In order to focus on potential distal cis-regulatory elements, we excluded accessible regions within 1kb upstream or 0.5 kb downstream of transcription start sites. This analysis yielded 33,679 distal accessible elements, with 5,736 elements over-enriched in chondrocytes and 8,955 elements under-enriched in chondrocytes (Figure 1B-C). As confirmation of sequence quality, the cartilage-specific R2 enhancer of the Collagen Type II alpha 1 a gene (*col2a1a*) is enriched in chondrocyte-specific elements (Figure 1-figure supplement 1A). De novo motif analysis of the top 2000 chondrocyte-enriched regions using HOMER recovered motifs for Sox, Fox, Nfat, Zfx and Nkx transcription factor families (Figure 1D; Supplementary file 1A). Sox9 ChIP-Seq of mouse chondrocytes had previously revealed Nfat and Fox motifs as the second and third most co-enriched with Sox motifs (Ohba et al., 2015), and the Nkx motif might reflect the role of Nkx3.2 in cartilage differentiation (Provot et al., 2006). Consensus sequences for Sox, Fox, and Nfat motifs were highly similar between zebrafish and mouse (Figure 1-figure supplement 1B), despite our zebrafish analysis focusing on all accessible regions in facial chondrocytes and mouse analysis focusing on only Sox9-bound regions in rib chondrocytes. These striking motif similarities indicate strong conservation of the cartilage gene regulatory network between fish and mammals, and between the face and rib, and strongly suggest that many of the identified chondrocyte-specific accessible regions in zebrafish likely function as chondrocyte enhancers. In addition, Gene Ontology (GO) analysis of the nearest genes to the chondrocyte-enriched elements revealed cartilage development as one of the top 6 associated terms (Figure 1E). We also recovered terms for neural crest cell migration and dorsal/ventral pattern formation, likely reflecting retention of enhancer accessibility linked to the neural crest origins and later dorsoventral arch patterning of the precursors of facial cartilage.

**Figure 1.**
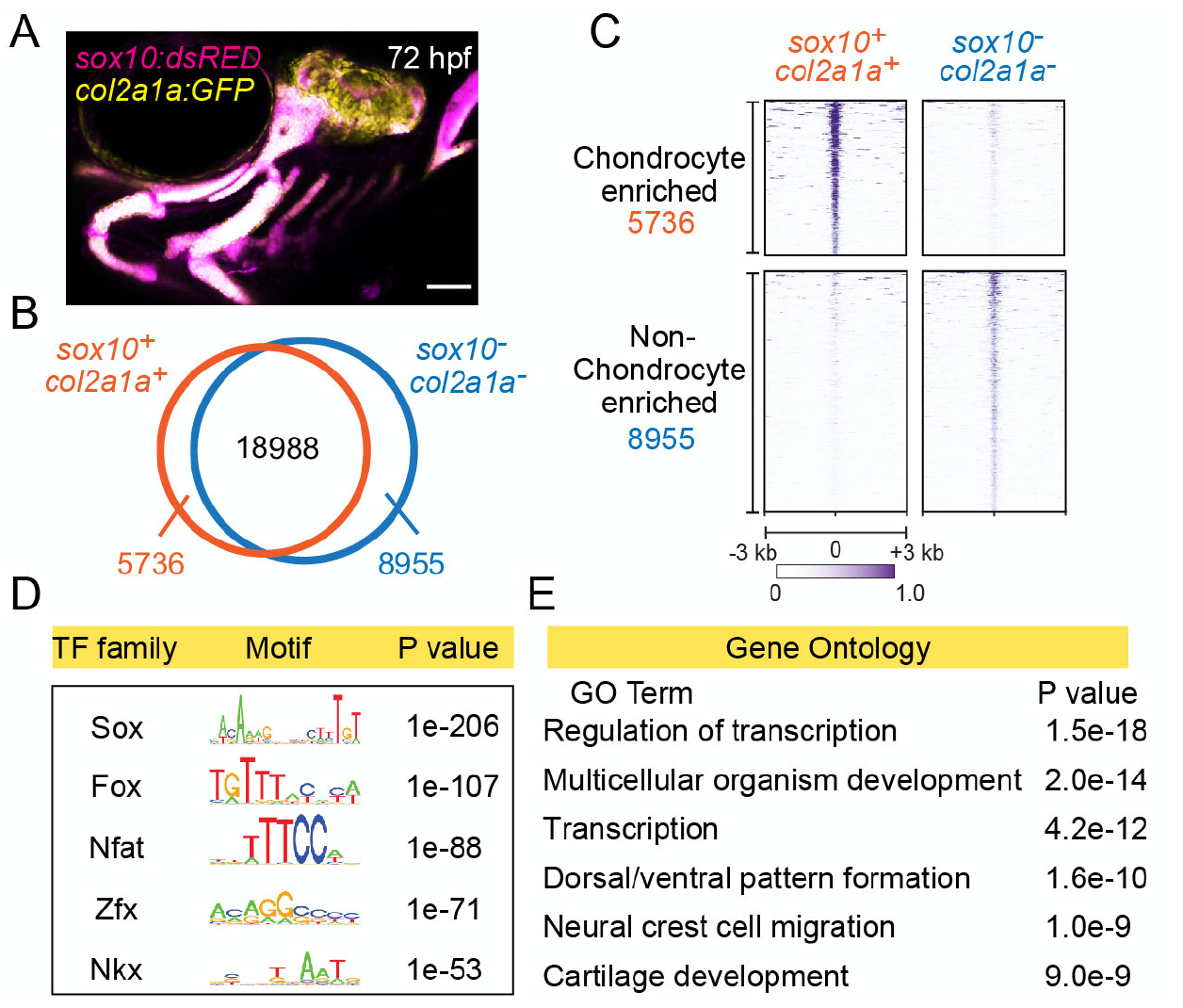
Chromatin accessibility landscape of facial chondrocytes. (A) Confocal image of facial cartilages expressing *col2a1a:GFP* and *sox10:Dsred* at 72 hpf. Lateral view with anterior to left. Scale bar = 100 μm. (B) Venn diagram indicating distal elements with open chromatin accessibility in *col2a1a:GFP*+; *sox10:Dsred*+ versus *col2a1a:GFP*-; *sox10:Dsred*-cells. (C) Peak intensity plots (Heatmap) of μATACseq show differentially enriched open chromatin regions in double-positive versus double-negative cells. (D) The top 5 transcription factor (TF) motifs recovered from the top 2000 μATACseq peaks enriched in chondrocytes (after removing redundant motifs). (E) GO analysis of nearest neighbor genes of μATACseq peaks enriched in chondrocytes.

### Progressive establishment of cartilage-associated chromatin accessibility

We next investigated when cartilage-associated chromatin accessibility is established in relation to CNCC development. CNCCs are first specified at ~10.5 hpf at the border of the neural keel, finish their migration into the pharyngeal arches by ~20 hpf, and then show the first histological signs of cartilage development in the jaw at ~52 hpf (Schilling and Kimmel, 1997). CNCC-derived arch ectomesenchyme cells can be uniquely identified by co-expression of *fli1a:GFP* and *sox10:Dsred* transgenes at 36 hpf and 48 hpf (Askary et al., 2017), stages just prior to cartilage differentiation (Figure 2A). We performed μATACseq on *fli1a:GFP*+; *sox10:Dsred*+ cells after FACS and analyzed chondrocyte-specific accessible elements from our 72 hpf dataset for their accessibilities at 36 and 48 hpf (Figure 2B). Of the 5,736 elements enriched in chondrocytes at 72 hpf, only 6% (356) had peak accessibility at 36 hpf with no further increase in accessibility by 48 hpf (“Group I”). In contrast, 48% (2,741) displayed increased accessibility between 36 and 48 hpf (“Group II”), and 46% (2,639) between 48 and 72 hpf (“Group III”). For Group I, de novo motif enrichment revealed predicted binding sites for members of the Nfat, Fox, Lhx, Nr2f, Meis and Pax families, and significant GO terms included neural crest migration, cell migration in general, and dorsal/ventral pattern formation (Figure 2C,D). Combined with the known involvement of Foxd3 (Montero-Balaguer et al., 2006; Stewart et al., 2006), Lhx6/8 (Denaxa et al., 2009), Nr2f1/2/5 (Barske et al., 2018), Meis2 (Machon et al., 2015) and Pax9 (Nakatomi et al., 2010) in CNCC specification, migration, and dorsal-ventral arch patterning, many Group I elements likely represent retention of cis-regulatory elements involved in the earlier specification, migration, and regional patterning of CNCCs. Group II and Group III elements share many common predicted transcription factor binding motifs, including Sox dimer, Fox, Nfat, and Ap1 motifs previously described for mouse cartilage (Figure 2E,G; Supplementary file 1B) (Ohba et al., 2015). An Nkx motif was recovered only for Group II (*p*=10^−58^, 36% of targets), an Egr motif was enriched for Group II (*p*=10^−65^, 55% of targets) versus Group III (*p*=10^−20^, 16% of targets), and a Tead motif was enriched for Group III (*p*=10^−63^, 36% of targets) versus Group II (*p*=10^−20^, 3% of targets). GO analysis for linked genes also revealed terms related to skeletal system development (Group II) and cartilage development (Group III), as well as more general terms such as transcription and organism development (Figure 2F,H). We therefore conclude that the majority of chondrocyte-specific elements gain accessibility after pharyngeal arch formation, and that transcription factor binding motifs change during cartilage differentiation. For example, enrichment of Nkx motifs in Group II elements might reflect the role of Nkx3.2 in limiting chondrocyte maturation (Provot et al., 2006) and promoting joint formation (Miller et al., 2003), while the preferential enrichment of the Tead motif, which is linked to growth-associated Hippo signaling (Ota and Sasaki, 2008), in Group III elements might reflect the later proliferative expansion of chondrocytes.

**Figure 2.**
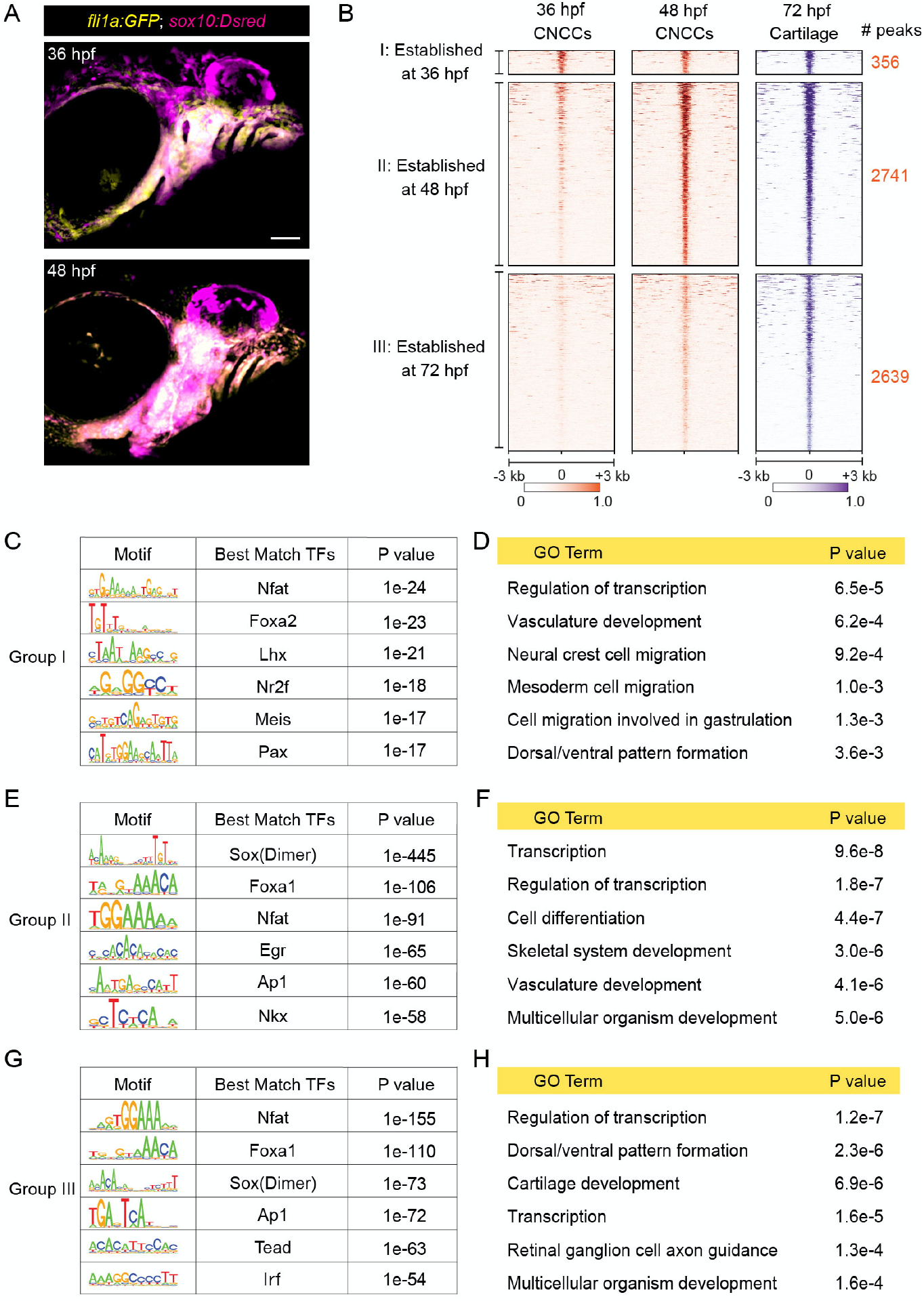
Dynamics of chromatin accessibility across facial chondrogenesis. (A) Confocal images of CNCCs expressing *fli1a:GFP* and *sox10:Dsred* at 36 and 48 hpf. Lateral view with anterior to left. Scale bar = 100 μm. (B) Peak intensity plots of cartilage-accessible distal elements shown for chondrocytes at 72 hpf and CNCCs at 36 and 48 hpf. Chondrocyte accessible elements are pooled into three categories based on dynamics of chromatin accessibility across stages. (C, E, G) De novo motif enrichment recovered by Homer analysis among the three categories. Top 6 motifs are shown with associated P values after removing redundant motifs. (D, F, G) GO term analysis among the three categories.

### Requirement of Foxc1 for chromatin accessibility at a subset of cartilage elements

We had previously found that Foxc1 genes are essential for cartilage development in the upper face (Xu et al., 2018), and both our μATACseq analysis of zebrafish chondrocytes and published Sox9 ChIP-seq analysis in mouse (Ohba et al., 2015) reveals co-enrichment of Sox and Fox motifs in accessible regions near known cartilage genes. In zebrafish Foxc1 (*foxc1a−/−; foxc1b−/−*) mutants, cartilages of the upper/dorsal face fail to develop (Xu et al., 2018). In order to isolate the dorsal CNCC precursors affected in Foxc1 mutants, we used a *pou3f3b:Gal4; UAS:nlsGFP* (*pou3f3b>GFP*) dorsal CNCC transgenic line (#Barske et al., 2020, PNAS, in press) along with the pan-CNCC *sox10:Dsred* transgenic line. Whereas *pou3f3b>GFP+; sox10:Dsred+* CNCCs were present in Foxc1 mutant pharyngeal arches at 48 hpf, many fewer differentiated into chondrocytes by 6 days post-fertilization (dpf) compared to sibling controls (Figure 3A-D). We therefore performed μATACseq on *pou3f3b>GFP+; sox10:Dsred+* cells after FACS from Foxc1 mutants and controls at 36 and 48 hpf. A comparison of the 15,781 accessible regions in dorsal CNCCs (*pou3f3b>GFP+; sox10:Dsred+*) and pan-CNCCs (*fli1a:GFP+; sox10:Dsred+*) at 36 hpf revealed 79% with similar accessibility, 11% with greater accessibility in dorsal CNCCs, and 10% with greater accessibility in pan-CNCCs (Figure 3-figure supplement 1A). Of 22,323 elements at 48 hpf, 72% were similarly accessible, 10% more accessible in dorsal CNCCs, and 18% more accessible in pan-CNCCs (Figure 3-figure supplement 1B). A comparison of the 5,736 regions with specific accessibility in cartilage at 72 hpf revealed high correlation between dorsal- and pan-CNCCs at 36 hpf, with 96% displaying similar accessibility (r=0.92, Figure 3-figure supplement 2A,C). By 48 hpf, however, we observed notable differences between accessibility of cartilage-specific elements, with 43% displaying greater accessibility in pan-CNCCs and only 0.2% displaying greater accessibility in dorsal CNCCs (r=0.71, Figure 3-figure supplement 2B,D). The decreased accessibility of cartilage-specific elements in dorsal CNCCs at 48 hpf supports previous studies that dorsal chondrocytes develop later than other chondrocytes in the zebrafish face (Schilling and Kimmel, 1997; Barske et al., 2016).

**Figure 3.**
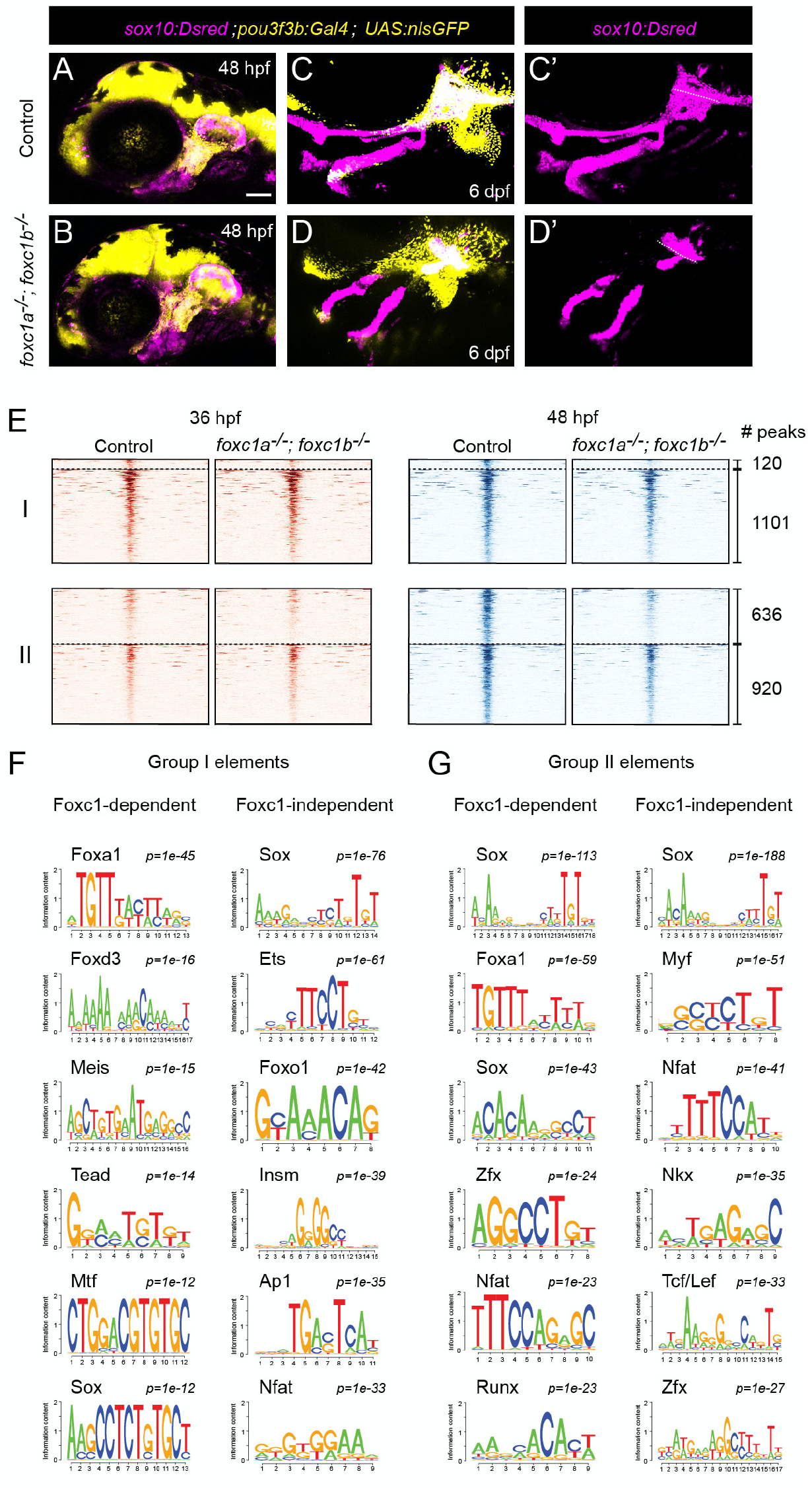
Foxc1 dependency of facial chondrocyte chromatin accessibility. (A, B) Confocal images show dorsal CNCCs of the first two arches labeled by *sox10:Dsred* and *pou3f3b:Gal4; UAS:nlsGFP* in control and *foxc1a*^−/−^; *foxc1b*^−/−^ mutant embryos at 48 hpf. Scale bar = 100μm. (C, D) Confocal images show loss of dorsal cartilages in *foxc1a*^−/−^; *foxc1b*^−/−^ mutant embryos at 6 dpf. *sox10:Dsred*+ cartilages are seen in single channels in C’ and D’, with dashed lines highlighting boundaries of dorsal and otic cartilage. (E) Peak intensity plots of Group I and Group II elements in control and *foxc1a*^−/−^; *foxc1b*^−/−^ mutant embryos. Peaks above the dashed lines are reduced in mutants. (F, G) De novo motif enrichment of Foxc1-dependent and Foxc1-independent Group I and Group II elements. Top 6 motifs are shown with associated P values after removing redundant motifs.

Analysis of cartilage-associated elements in Foxc1 mutants revealed that 10% (120/1,221) of elements that have established peak accessibility by 36 hpf (i.e. Group I) and 41% (636/1,556) of elements that increase accessibility between 36 and 48 hpf (i.e. Group II) had reduced accessibility in Foxc1 mutants (Figure 3E). De novo motif analysis of Foxc1-dependent and Foxc1-independent elements in Group I showed enrichment of Sox (*p*=10^−12^, *p*=10^−76^), Tead (*p*=10^−14^, *p*=10^−20^) Mtf (*p*=10^−12^, *p*=10^−22^) and Ets (*p*=10^−12^, *p*=10^−61^) motifs in both (Figure 3F, Supplementary file 1C). Whereas Fox motifs were uncovered in both, a closer analysis revealed enrichment of Foxa2 and Foxd3 motifs only in Foxc1-dependent elements (*p*=10^−45^, 38% of targets; *p*=10^−16^, 18% of targets), and a Foxo1 motif only in Foxc1-independent elements (*p*=10^−42^, 49% of targets). Similarly in Group II elements, a Foxa1 motif was enriched only in Foxc1-dependent elements (*p*=10^−59^, 38% of targets), and Foxh1 and Foxp1 motifs only in Foxc1-independent elements (*p*=10^−22^, 22% of targets; *p*=10^−19^, 7% of targets) (Figure 3G, Supplementary file 1C). Although Foxc1 motifs were not in the database used for motif predictions, the selective presence of Foxa1/2 motifs in Foxc1-dependent elements is consistent with previous reports that the consensus sequence for Foxc1-bound peaks is nearly identical to Foxa1/2 motifs (Wang et al., 2016). Sox motifs were similarly enriched in both Group I and Group II Foxc1-dependent and -independent elements, and Nfat and Zfx motifs in Group II Foxc1-dependent and -independent elements. An Insm (*p*=10^−39^, 30% of targets) motif was uncovered only in Foxc1-independent Group I elements, and an Nkx motif (*p*=10^−35^, 52% of targets) only in Foxc1-independent Group II elements. These findings suggest that Foxc1-dependent and -independent cartilage elements are commonly bound by Sox9 but likely differ in co-binding by Foxc1 and additional co-factors. For example, the presence of the Nkx motif only in Foxc1-independent elements suggests that it could be an alternative co-factor for Sox9, in line with the known roles for Nkx3.2 in chondrocyte biology (Provot et al., 2006).

### Validation of Foxc1-dependent cartilage enhancers in zebrafish transgenesis assays

To verify whether Foxc1-dependent cartilage elements identified by μATACseq are chondrogenic enhancers, we tested the ability of individual elements in combination with an E1b minimal promoter to drive cartilage expression of green fluorescent protein (GFP) in zebrafish transgenic assays (Figure 4-figure supplement 1A). We tested 22 Foxc1-dependent elements near 15 different genes, which included elements linked to genes with known cartilage function (*ucmab, matn4, matn1, lect1, epyc, col9a1a, col9a3, sox10, acana, foxa3, mia*) and others with unknown cartilage function (*si:dkey33i1l.4, gas1b, lefty2, slc35d1a*) (Figure 4A; Figure 4-figure supplement 1; Supplementary file 2). We observed that 59% (13/22) of elements drove GFP expression in facial chondrocytes at 6 dpf. These included intronic elements within *sox10, lect1, col9a3, col9a1a* and *slc35d1a*; distal 5’ elements near *ucmab, epyc, mia, acana*, and *matn4*; a distal 3’ element near *gas1b*; and a promoter-associated element for *lect1*. All tested elements are from Group II, i.e. increasing accessibility between 36 and 48 hpf, and we confirmed cartilage expression in independent stable transgenic lines in 9 cases (Figure 4B; Figure 4-figure supplement 1; Supplementary file 2). Whereas an element in the first intron of *sox10* drove uniform cartilage-specific expression, most elements drove expression in specific sub-regions or differentiation stages of facial chondrocytes. Whereas a distal 5’ element of *ucmab* drove expression in chondrocytes of multiple joints in the zebrafish head, an intronic element of *lect1* drove chondrocyte expression only in the jaw joint and hyomandibular-otic connection, a promoter-associated element of *lect1* drove expression more strongly in the hyoid joint and hyomandibular-symplectic connection (though also more broadly in chondrocytes), and a distal 5’ element of *acana* drove restricted chondrocyte expression at the hyoid joint. Reciprocally, elements associated with *col9a3, epyc, mia*, and *col9a1a* were expressed in chondrocytes but generally excluded from joint regions (particularly apparent at the hyoid joint and hyomandibular-symplectic connection). Further, we found that three enhancer transgenes with diverse expression patterns (broad *sox10*, joint-restricted *ucmab*, and joint-excluded *epyc* enhancers) all displayed reduced activity specifically in the dorsal cartilage regions affected in Foxc1 mutants (Figure 4C). We also tested four Foxc1-independent elements and found that two drove cartilage expression (distal 5’ elements of *gas1b* and *matn4*), one drove ligament expression (distal 5’ element of *sparc*), and one had no activity (promoter-associated element of *sox10*) (Figure 4-figure supplement 1). Thus, the majority of accessible elements tested had chondrocyte activity, and Foxc1-dependent and -independent elements appear equally capable of driving cartilage expression. Moreover, the activity of different enhancers in diverse locations and stages of chondrocyte maturation indicates that global cartilage expression patterns are achieved in part through the summation of enhancers with more restricted activity.

**Figure 4.**
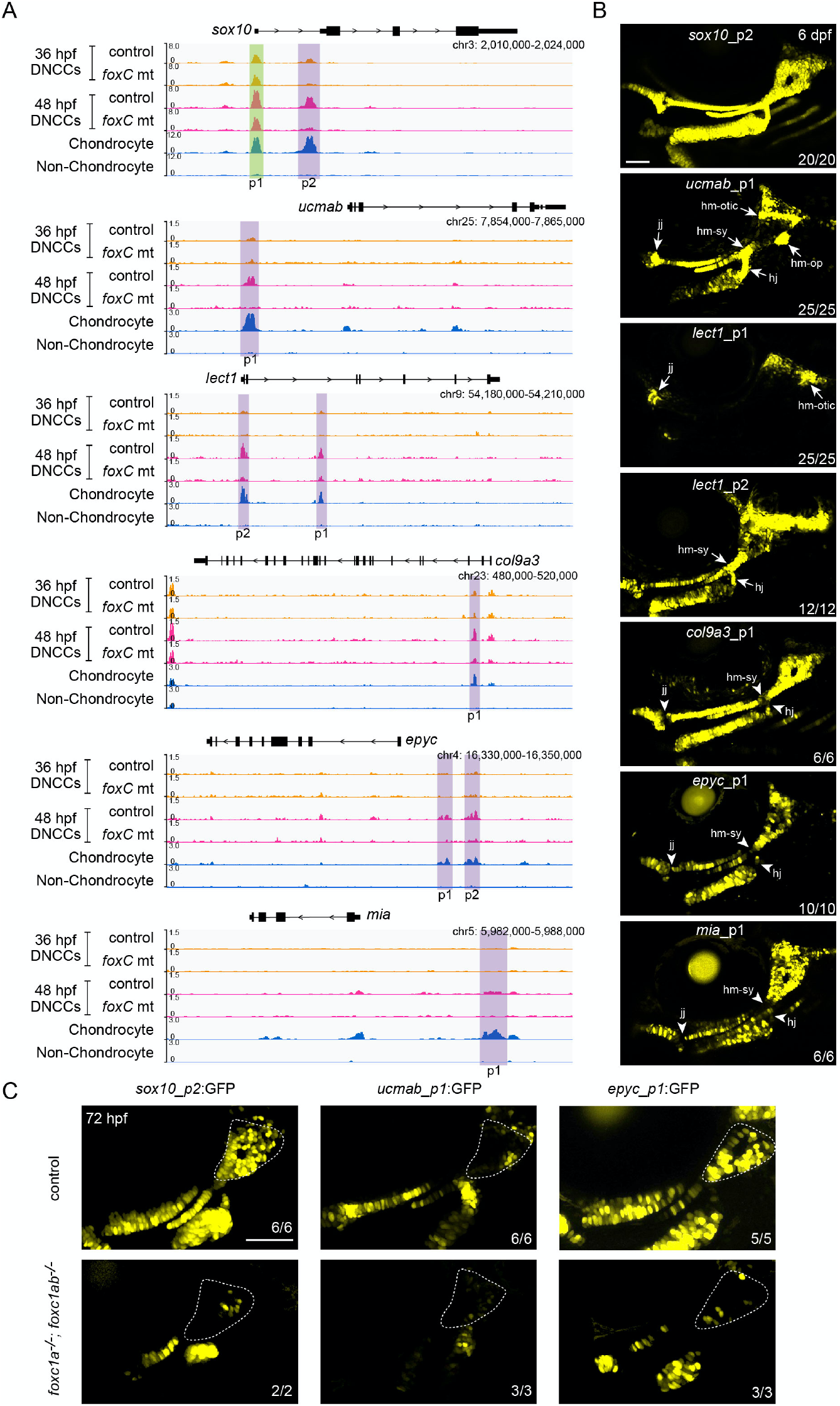
In vivo validation of Foxc1-dependent cartilage enhancers. (A) Snapshots of genomic regions (genes and GRCz10 coordinates listed) for enhancer testing. Peaks (p) tested are shown, with Foxc1-dependent regions in purple and a Foxc1-independent element in green. μATACseq reads are shown in each row for the experiments indicated, with chondrocyte and non-chondrocyte peaks from 72 hpf embryos. DNCCs, dorsal CNCCs. (B) GFP expression driven by the indicated peaks in stable transgenic zebrafish at 6 dpf. Confocal projections of the cartilages of the first two arches are shown in lateral view with anterior to the left. Arrows indicate enriched expression at joint regions, and arrowheads denote relative lack of expression. jj, jaw joint. hj, hyoid joint. hm-sy, hyomandibular-symplectic junction. hm-otic, hyomandibular-otic junction. hm-op, hyomandibular-opercular joint. (C) Confocal projections show selective loss of *sox10_p2:EGFP, ucmab_p1:EGFP*, and *epyc_p1:EGFP* transgene expression in the dorsal cartilage domains (dashed outlines) of *foxc1a*^−/−^; *foxc1b*^−/−^ mutants at 72 hpf. Numbers indicate proportion of embryos in which the displayed patterns were observed. Scale bars = 100 μm.

## Conclusion

Our findings indicate that Foxc1 function is required for the accessibility of close to half of chondrocyte enhancers in the zebrafish face. Given expression and function of Foxc1 in diverse cartilages (e.g. limb, rib, tracheal) in mouse, it seems likely that Foxc1 has a similar function in chondrocyte enhancer accessibility throughout the body. It is unclear why only a minority of enhancers appears to require Foxc1 activity. It is possible that other members of the Fox family compensate, such as Foxf1/2 in the facial midline (Xu et al., 2018) and Foxa2/3 during later hypertrophic maturation (Ionescu et al., 2012). Alternatively, there may be other co-factors that cooperate with Sox9 for chondrocyte enhancer accessibility and activation, such as Nkx3.2. We did not detect any obvious differences in the types of enhancer-proximal genes or the patterns and stages associated with Foxc1-dependent versus-independent enhancer activity. Further work will also be needed to understand the mechanism by which Foxc1 promotes chondrocyte enhancer accessibility. Foxc1 lacks a chromatin modifying domain (Yoshida et al., 2015) and therefore would need to interact with a co-factor to directly open chromatin. Foxc1 could also act to maintain open chromatin, as shown for Foxa1 in the liver (Reizel et al., 2020). Given that both Foxc1 and Sox9 have expression in many tissues outside the skeletal system, it will also be important to determine whether additional factors help to further restrict their activity to chondrocyte enhancers within skeletogenic mesenchyme.

## Methods

### Zebrafish lines

The Institutional Animal Care and Use Committee of the University of Southern California approved all experiments on zebrafish (*Danio rerio*) (Protocol #10885). Existing mutant and transgenic lines used in this study include *foxc1a^el542^* and *foxc1b^el620^* (Xu et al., 2018); Tg(*sox10:Dsred*)^el110^ and *Tg*(*fli1a:EGFP*)^y1^ (Askary et al., 2017); *Tg*(*col2a1aBAC:GFP*) (Paul et al., 2016); and *Tg*(*UAS:nlsGFP;α-crystallin:Cerulean*)^*el609*^ *and pou3f3b^Gal4ff-el79^* (#Barske et al., 2020, PNAS, in press). For enhancer transgenic lines, we synthesized accessible elements with flanking attB4 and attB1 sequences using IDT gBlocks and cloned these into pDONR-P4-P1R using the Gateway Tol2kit (Invitrogen) to create p5E enhancer constructs (Kwan et al., 2007). We then combined p5E constructs with pME-E1b-GFP, p3E-polyA, and pDestTol2AB2 using LR clonase. Final DNA constructs were microinjected with transposase RNA (30 ng/μl each) into 1cell stage zebrafish embryos. In most cases, multiple independent stable alleles per construct were analyzed in the F1 generation (Supplementary file 2).

### Confocal imaging

Transgenic embryos were imaged with a Zeiss LSM800 confocal microscope. Maximum intensity projections are shown for stable transgenic lines and single representative sections are shown for injected embryos. Image levels were modified consistently across samples in Adobe Photoshop CS6.

### μATACseq

Wild-type embryos double-positive for *fli1a:EGFP* and *sox10:Dsred* (36 hpf and 48 hpf), or *col2a1a:GFP* and *sox10:Dserd* (72 hpf), were sorted under a fluorescent dissecting microscope (Leica M165FC) before dissociation. For Foxc1 mutant analysis, we performed incrosses of *pou3f3b:Gal4*+/−; *UAS:nls-GFP*+/−; *sox10:Dsred*+/−; *foxc1a*+/−; *foxc1b*+/− fish and then selected for GFP+/Dsred+ embryos on a fluorescent dissecting microscope. Genotyping was then performed on tail lysates collected from individual embryos at 27 hpf. We then pooled *foxc1a*−/−; *foxc1b*−/− double mutants and separate sibling controls (*foxc1a*+/−; *foxc1b*+/+ and *foxc1a*+/+; *foxc1b*+/+ embryos) for FACS. To facilitate embryo collection at the 36 hpf time point, embryos were moved at 27 hpf to an incubator set at 22°C to delay their development such that they reached 36 hpf the following morning. Cell dissociation and FACS were performed as previously described (Askary et al., 2017). Around 5000 cells of each sample were centrifuged at 500 g for 20 m at 4 °C, and the pellet was suspended with 20 μL of lysis buffer (10 mM Tris HCl (pH 7.4), 5 mM MgCl2, 10% DMF, 0.2% N-P40) by pipetting 6-10 times to release the nuclei without purification. The cell lysate was then mixed with 30 μL reaction buffer (10 mM Tris HCl (pH 7.4), 5 mM MgCl2, 10% DMF, and homemade Tn5 Transposase) by vortexing for 5 s. The reaction was incubated at 37°C for 20 minutes, followed by DNA purification using a Qiagen MiniElute kit. Purified DNA fragments were used to conduct μATACseq libraries as previously described (Buenrostro et al., 2013) and sequenced using the NextSeq 500 platform (Illumina) with a minimum of 50 million paired-reads/sample.

### Data analysis and statistics

The Encode analysis pipeline (https://github.com/ENCODE-DCC/chip-seq-pipeline) for ATACseq was used with small modifications. The raw reads were trimmed to 37 bp and aligned to zebrafish GRCz10 genome assembly by STAR aligner (Dobin et al. 2013). PCR duplicates, and the reads that aligned to “blacklist regions” (Amemiya et al. 2019) were removed, and then peaks were called by Model-based analysis of ChIP-Seq (MACS2) (Zhang et al. 2008) with *p*=0.01 cutoff and disabled dynamic lambda option (--nolambda) for individual replicates. Peaks from individual replicates were further filtered by IDR<0.01, and the overlapping peaks between replicates were used. Bigwig files were generated from dup-removed bam files with *bedtools* and *bedgraphtobigwig*, and normalization was based on total read numbers. Heatmaps were generated with DeepTools (Ramírez et al. 2016) based on the normalized bigwig signal files. Individual genomic loci were examined by IGV (Broad Institute). Differential analyses of μATACseq data was done with DESeq2 (Love et al. 2014). HOMER (Heinz et al. 2010) was used to identify de novo motifs. David 6.8 GO analysis was performed on the web interface (https://david.ncifcrf.gov/) based on nearest neighbor genes for all differentially accessible elements. Pearson correlation was calculated using cor() function in R.

## Acknowledgements

We thank Jeffrey Boyd at the USC Stem Cell Flow Cytometry Core for FACS, David Ruble at the CHLA Sequencing Core, the high-performance computing core at USC, and Megan Matsutani, Jennifer DeKoeyer Crump, and Mathi Thiruppathy for fish care. We dedicate this work to the late Bartosz Balczerski whose passion as a postdoc inspired this project.

## Author Contributions

P.X. and J.G.C. conceived of experiments. P.X., M.F, and P.F. conducted zebrafish experiments. P.X. and H.V.Y. performed μATACseq experiments, and with K.-C.T. conducted bioinformatics analysis. N.S. and J.G.C. obtained funding and oversaw project. P.X. and J.G.C. wrote the manuscript.

## Competing interests

The authors declare no competing interests.

## Materials & Correspondence

Requests for material should be directed to J. Gage Crump.

## Data availability

μATACseq data can be accessed at GEO with accession number GSE157575: visit https://www.ncbi.nlm.nih.gov/geo/query/acc.cgi?acc=GSE157575_;!!LIr3w8kk_Xxm!4o2V_xfIwp2fIZl1FVkwGRA5Q9CcHhvfeKfQYZftKgRy6JUHBaaAPO2btB-Xgs0o$ and enter token gzcxqyourvmdnuj into the box.

## SUPPLEMENTARY FIGURE LEGENDS

**Figure 1-figure supplement 1.**
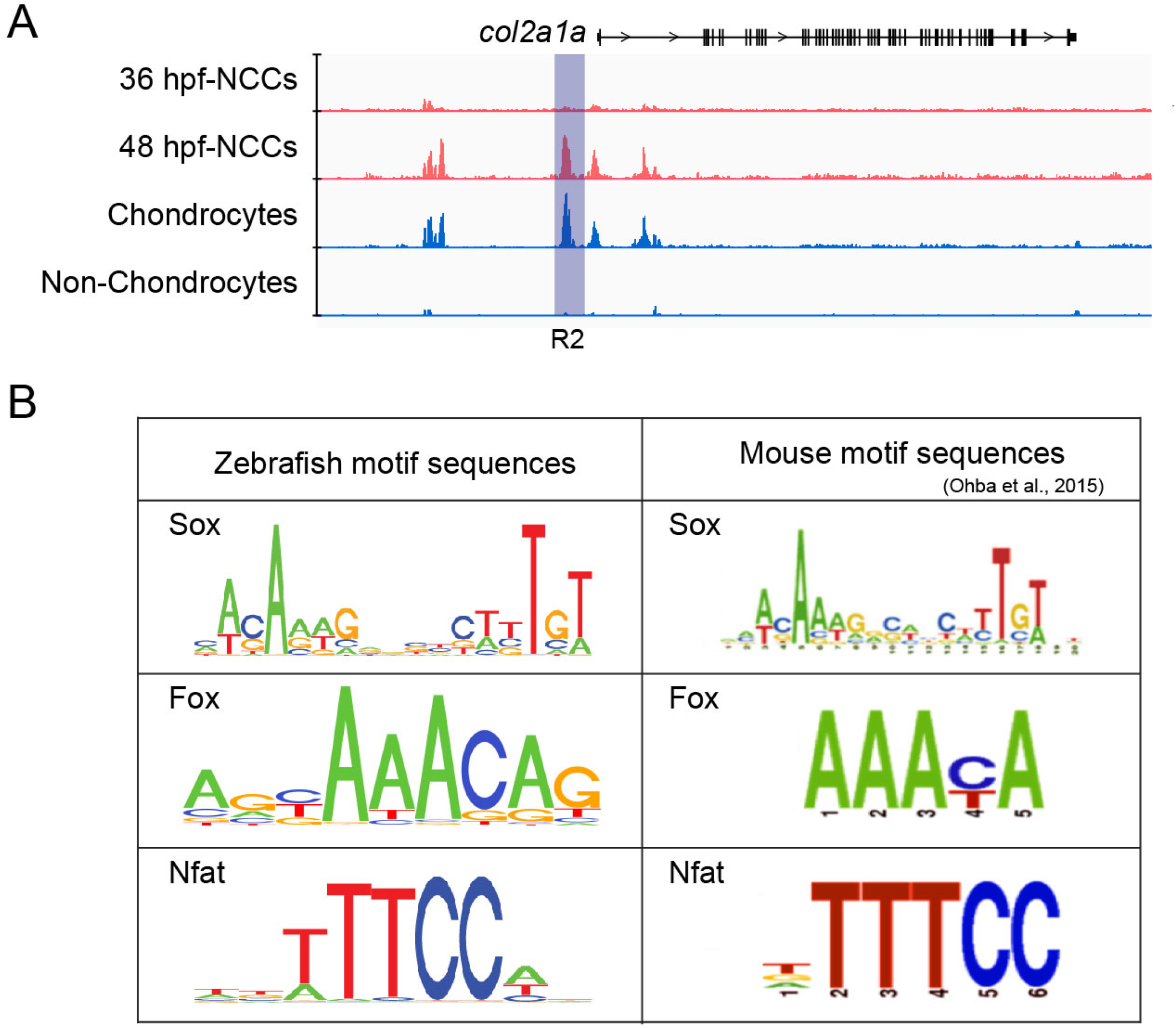
*col2a1a* enhancer and cartilage motif comparison between zebrafish and mouse. (A) Snapshot of the *col2a1a* locus showing μATACseq reads from the indicated experiments. Several regions become accessible in 48 hpf CNCCs and then remain open in chondrocytes at 72 hpf. Boxed region shows the published R2 cartilage enhancer. (B) Comparison of consensus motif sequences recovered from zebrafish μATACseq peaks enriched in facial chondrocytes (left) and mouse Sox9 ChIP-Seq peaks in rib chondrocytes (right).

**Figure 3-figure supplement 1.**
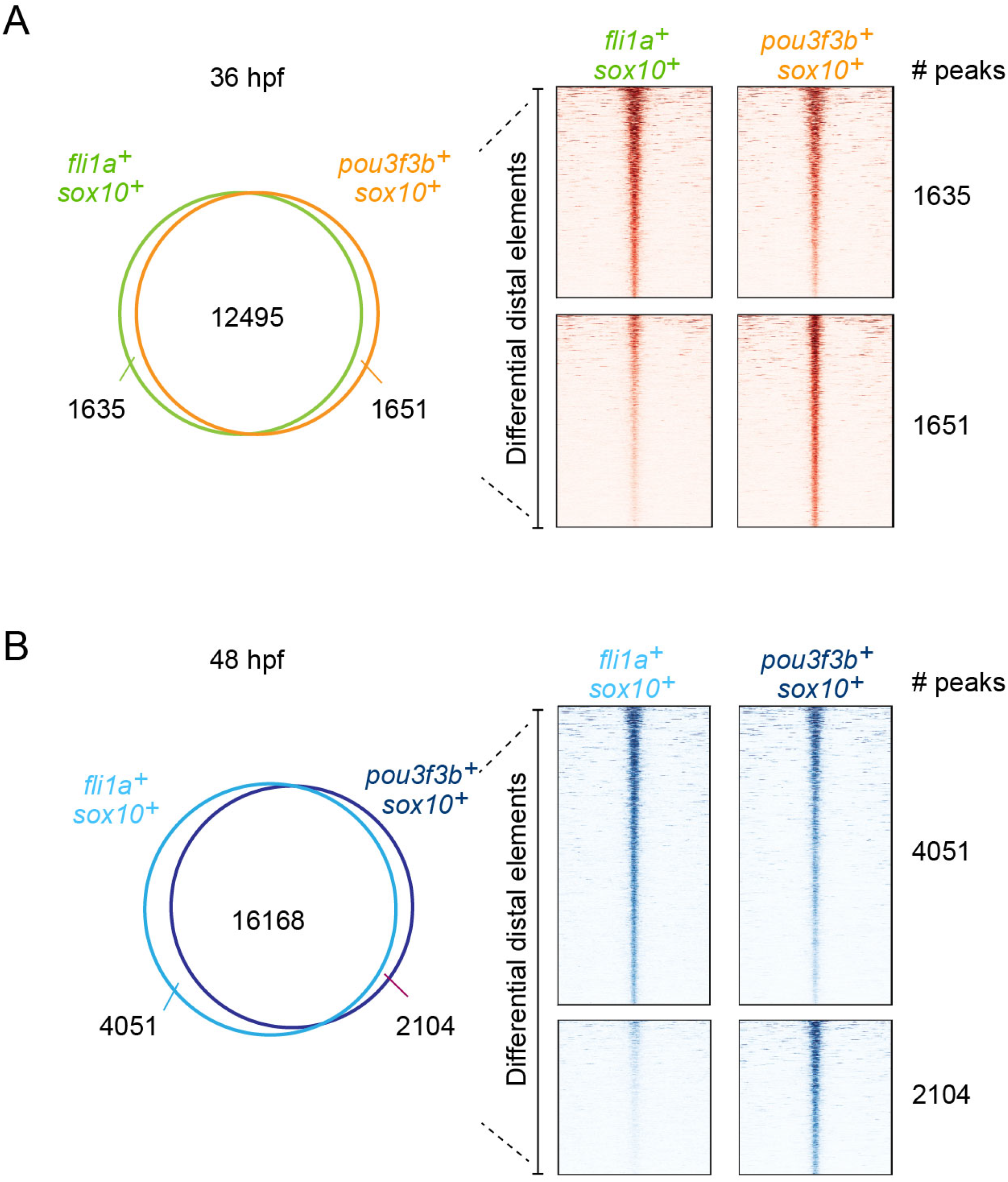
Comparison of accessible regions between pan- and dorsal CNCCs. Venn diagrams and heatmaps show distal accessible elements in μATACseq data from *fli1a:GFP*+; *sox10:Dsred*+ (pan-CNCCs) and *pou3f3b>GFP+; sox10:DsRed+* (dorsal CNCCs) at 36 hpf (A) and 48 hpf (B). Numbers list elements unique for each population or shared (center of circles).

**Figure 3-figure supplement 2.**
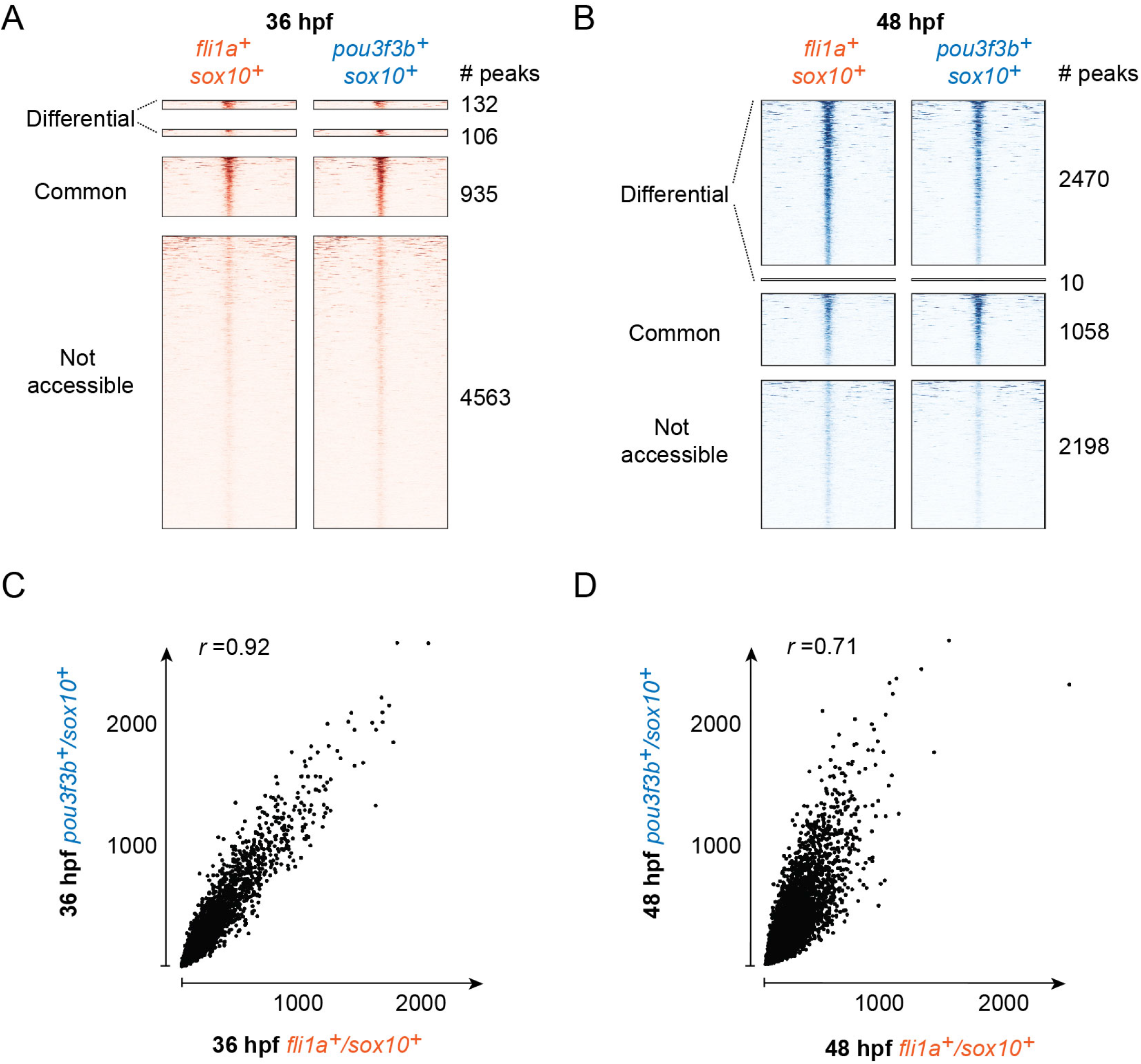
Accessibility of cartilage-enriched elements in earlier pan- and dorsal CNCCs. (A,B) Heatmaps of the 5,736 distal elements with accessibility in 72 hpf chondrocytes. Their accessibility is then plotted according to μATACseq data from *fli1a:GFP*+; *sox10:Dsred*+ (pan-CNCCs) and *pou3f3b>GFP+; sox10:Dsred+* (dorsal CNCCs) at 36 hpf (A) and 48 hpf (B). At 36 hpf, 132 peaks were more accessible in pan-CNCCs and 106 peaks more accessible in dorsal CNCCs. At 48 hpf, 2,470 peaks were more accessible in pan-CNCCs and 10 peaks more accessible in dorsal CNCCs. Peaks commonly or not accessible in both data sets are also shown. (C,D) Pearson correlation graphs of the behavior of cartilage-associated distal elements in pan-versus dorsal CNCCs at 36 and 48 hpf. x and y axes represent raw read counts from BAM files, and “r” is the Pearson correlation coefficient.

**Figure 4-figure supplement 1.**
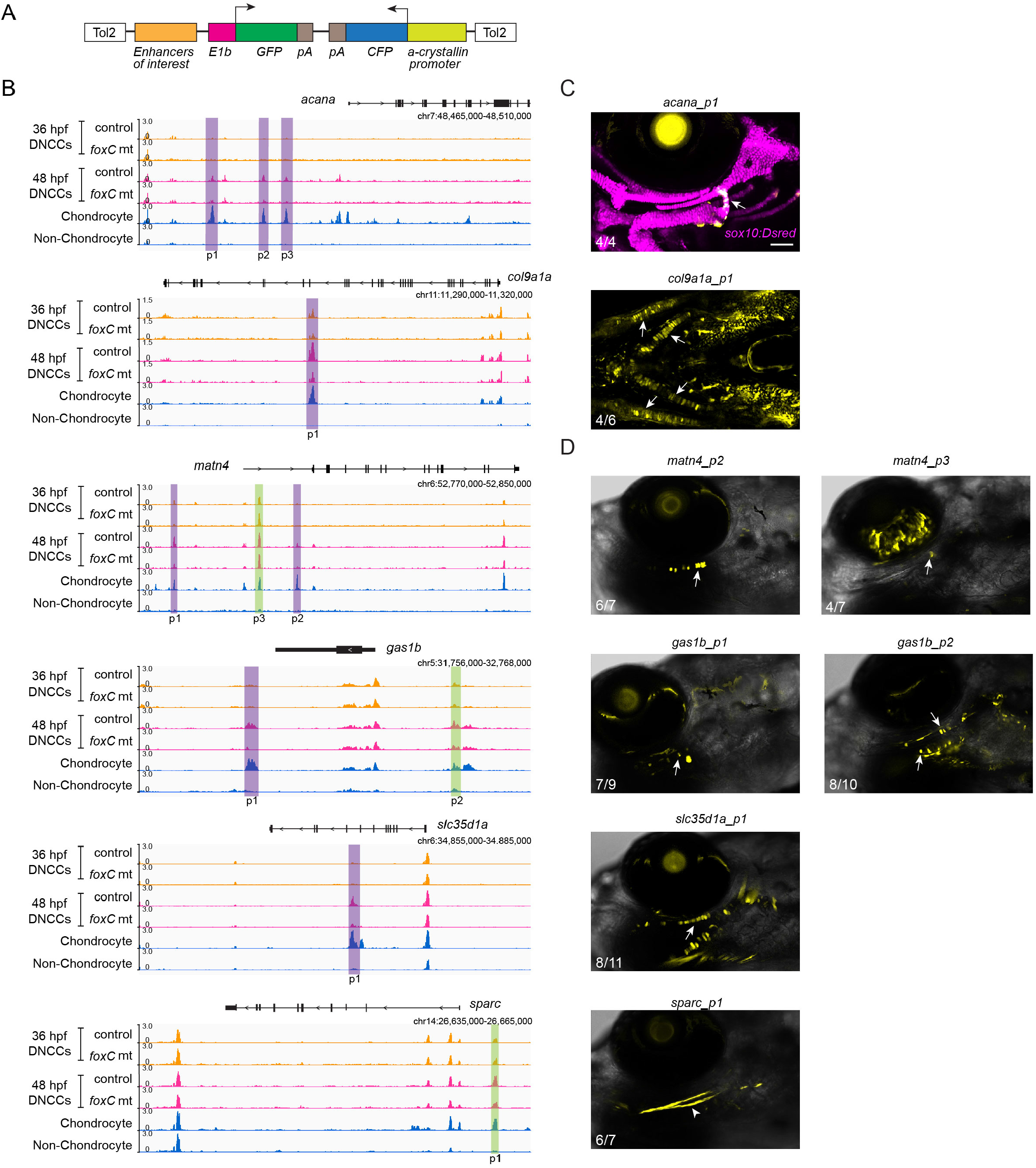
Additional in vivo validation of cartilage-enriched accessible elements. (A) Schematic of the reporter construct for enhancer testing. Tol2, transposase integration site. E1b, minimal promoter. GFP, green fluorescent protein. pA, polyadenylation sequence. CFP, cerulean fluorescent protein. (B) Snapshots of genomic regions (genes and GRCz10 coordinates listed) for enhancer testing. Peaks (p) tested are shown, with Foxc1-dependent regions in purple and Foxc1-independent elements in green. μATACseq reads are shown in each row for the experiments indicated, with chondrocyte and non-chondrocyte peaks from 72 hpf embryos. DNCCs, dorsal CNCCs. (C) GFP expression (yellow) driven by the indicated peaks in stable transgenic zebrafish at 6 dpf. Confocal projections of the cartilages of the first two arches are shown in lateral view for *acan_p1* and ventral view for *col9a1a_p1*. For *acan_p1, sox10:Dsred* labels all chondrocytes for reference in magenta, and arrow indicates hyoid joint expression. For *col9a1a_p1*, arrows indicate expression in the ventral Meckel’s and ceratohyal cartilages. (D) Confocal sections of representative embryos injected with enhancer constructs are shown in lateral view at 6 dpf. Mosaic expression in chondrocytes (yellow, arrows) is shown with DIC providing embryo context in white. For *sparc_p1*, interopercular ligament expression is denoted by arrowhead. Numbers indicate proportion of embryos in which the displayed patterns were observed. Scale bar = 100 μm.

